# Variants identify sarcomere inter-protein contacts distinguishing inheritable cardiac muscle diseases

**DOI:** 10.1101/2022.03.23.485392

**Authors:** Thomas P. Burghardt

## Abstract

Human ventriculum myosin (βmys) powers contraction sometimes while complexed with myosin binding protein C (MYBPC3) on the myosin thick filament. The latter regulates βmys activity through inter-protein contacts. Single nucleotide variants (SNVs) change protein sequence in βmys or MYBPC3. They cause inheritable heart disease. When a SNV modified domain locates to an inter-protein contact it affects complex coordination. Domains involved, one in βmys and the other in MYBPC3, form coordinated domains called co-domains. Co-domains are bilateral implying the potential for a shared impact from SNV modification in either domain suggesting their joint response to a common perturbation assigns location. Human population genetic divergence is the common systemic perturbation. A general contraction model with a neural/Bayes network design reveals SNV probabilities specifying correlations between domain members using 2D correlation genetics (2D-CG). It reveals co-domain locations in three common human heart diseases caused by SNVs, familial hypertrophic cardiomyopathy (FHC), dilated cardiomyopathy (DCM), and left ventricle non-compaction (LVN). Co-domain maps for DCM and LVN link MYBPC3 with two levels of myosin heads on the myosin thick filament surface implying these myosin dimers form the super-relaxed state (SRX). The FHC co-domain map involves just one myosin dimer implying the myosins do not form SRX. Comparing co-domain maps for FHC, DCM, and LVN phenotypes suggests SRX disruption involves a co-domain between MYBPC3 regulatory domain and the myosin regulatory light chain (RLC) N-terminus. The general contraction model scenarios, constructed from feed-forward neural networks, were explored with the purpose to understand how to interpret them mechanistically with basic natural language characteristics. These characteristics emerge from dependencies among inputs coded in hidden layer width and depth when they are deciphered using 2D-CG. In this application, the thick filament structural states emerge for FHC, DCM, and LVN phenotypes defining thick filament structural state joining the other standard characteristics of phenotype and pathogenicity. Emergent natural language interpretations for general network contraction models are on the horizon.

## INTRODUCTION

Muscle proteins assembled in the sarcomere produce contraction force and displacement by their coordinated action. In human ventriculum the cardiac myosin motor (βmys) repetitively converts ATP free energy into work while sometimes complexed with myosin binding protein C (MYBPC3). The latter regulates βmys and muscle activity through multiple inter-protein contacts between domain sub-structures of βmys and MYBPC3 [2–6]. Single nucleotide variants (SNVs) causing inheritable heart disease frequently target the βmys/MYBPC3 complex causing structural modification to the domains containing them [7]. When a SNV modified domain locates to an inter-protein contact it affects complex coordination. Domains involved, one in βmys and the other in MYBPC3, implies SNV modification probabilities for either domain will correlate jointly with a common systemic perturbation. Identifying their joint correlation assigns their location. βmys and MYBPC3 SNVs have pathogenicities that correlate with human populations [8] indicating that cardiac muscle physiology integrates βmys/MYBPC3 functionality with the wider system involving genetic background [9]. Population genetic divergence provides the common systemic perturbation assigning co-domains in the βmys/MYBPC3 complex [10].

Human population genetic divergence from a single origin in East Africa is attributed to the serial founder effect addressing migration [11], colonization, and exchange between geographically near populations [12]. It rationalizes the observed linear genetic divergence decrease with human migration distance over the earth’s surface that is likewise detected in a SNV database independently characterizing worldwide genetic variation [13]. It shows that migration distance is a good proxy for genetic divergence variation providing the means to use it in an ordered sequence of human populations to perturb SNV probability. Genetic divergence over human populations couples SNV probability across co-domains detected by cross-correlation using 2-dimensional correlation genetics (2D-CG) [10]. 2D-CG is a 2D correlation spectroscopy analog [14] wherein βmys and MYBPC3 functional domains replace the two spectral frequencies under surveillance, SNV probability products mimic resonance absorption intensity, and human population genetic differentiation provides the linear perturbation.

SNVs in βmys or MYBPC3 affect human populations everywhere. Public databases inform their impact on the human genetic coding with specific SNV data including four input parameters that are known with relative certainty: sequence position, residue substitution, human population, and frequency. And two output parameters sometimes unknown or unreported: phenotype and pathogenicity. This 6-dimensional data point (6ddp) parameter set describes functionality of a SNV modified human ventricular sarcomere. It is a system captured here in directed acyclic graphs (DAGs) relating input molecular level parameters to outputs describing the state of human health (**Figure 1**). The DAG represents a general contraction model interpreting molecular data as a systemic outcome. It is formulated for quantitation by a feed-forward neural network with subsets of completed 6ddps, where input and output parameters are known, forming the validation dataset. Incomplete 6ddps, where one or both output parameters are unknown, are completed using the optimized neural network. The full data set, now complete, is interpreted probabilistically with a discrete Bayes network also based on the DAG to give the SNV probability for a functional domain location given phenotype and human population. It supplies the information needed to identify phenotype specific co-domains in the βmys/MYBPC3 complex using 2D-CG. The approach, described earlier for application to the βmys/MYBPC3 complex [10], now includes improvements to the neural network optimization protocol giving a large accuracy boost.

**Figure 1.**
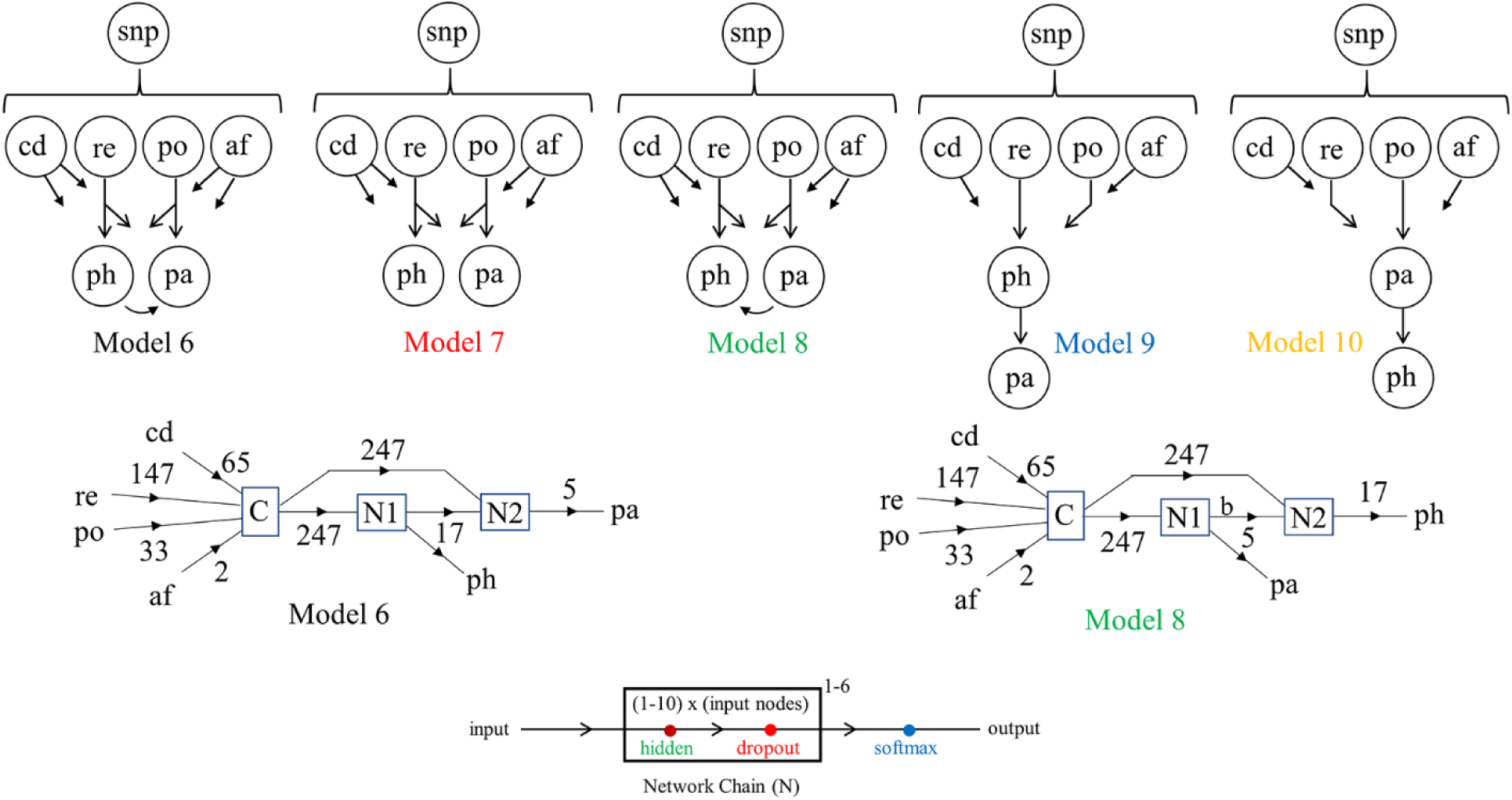
Directed acyclic graph (DAG) general contraction models (top row) for the βmys/MYBPC3 complex and their neural network representation (middle and bottom rows). Model 6, introduced previously, and 4 others systematically permute the path of influence connecting input with output parameters. Input parameters are functional protein domains (cd), residue replacement (re), human population (po), and SNV allele frequency (af). Output parameters are disease phenotype (ph) and pathogenicity (pa). Arrows show the direction of influence with no feedback. Neural networks (middle) indicate the input parameters concatenated at the C-module. Numbers over arrows in the diagrams in the middle row indicate parameter number (e.g., 65 domains for the βmys/MYBPC3 complex at cd) that are input to Network Chains (N) depicted with more detail in the bottom row. Input nodes to each network chain is equal to the total parameters input. Network chains have 1-6 hidden layers (indicated by the 1-6 superscript above the box in the net chain diagram) with hidden layer nodes numbering 1-10 times the number of input nodes. The dropout layer mitigates overfitting (along with another measure described in Methods) and the softmax layer converts the final output into digital form.

In addition, focus narrows onto three frequent heart disease phenotypes caused by SNV’s in the βmys/MYBPC3 complex, familial hypertrophic cardiomyopathy (FHC), dilated cardiomyopathy (DCM), and left ventricle non-compaction (LVN). FHC is the most common inheritable heart disease in the United States and is the leading cause of sudden cardiac death in the young and heart failure in the elderly. FHC involves hypercontractility characterized by hypertrophy of the cardiac muscle tissue and less efficient blood flow production [15]. DCM has a thinning cardiac muscle in one or more heart chambers. It is a progressive disease as chambers enlarge and weaken eventually becoming unable to produce sufficient blood flow. LVN affected cardiac muscle has thick left ventricle walls with a spongy appearance. Tissue fails to develop normally and causes progressive cardiac dysfunction in adults [16]. Their co-domain maps identify each phenotype by a bi-molecular interaction footprint reflecting their divergent molecular mechanisms.

Interpreting general contraction model scenarios found for the diseased systems with basic natural language structural characteristics is a wider project goal also explored. It is shown that structural characteristics emerge from dependencies among the 6ddps that are deciphered by 2D-CG. In this application 2D-CG informs the thick filament structural states distinguishing FHC, DCM, and LVN phenotypes defining a basic structural output joining phenotype and pathogenicity in an appended SNV database. Emergent natural language interpretations for general network contraction models are on the horizon.

## RESULTS

### Neural Network Models for Contraction

Neural networks following DAG designs in **Figure 1** predict phenotype and pathogenicity for comparison with values in the fulfilled 6ddps to evaluate design accuracy as cardiac ventriculum contraction models. Correct prediction fraction (CPF), given by the number of correct predictions divided by the sum of phenotype and pathogenicity data points in the fulfilled 6ddps (Methods), quantitates comparison. Perfect model equivalence to data has CPF equal to 1.

**Figure 2** shows CPF probability for each model scenario optimized by changing hidden layer depth and width in network chains (N1 and N2). Optimization implies searching through 1-6 hidden layers, and the (1-10) x (input node count) hidden layer widths for the 5 DAG inspired models. Each optimization run is called a trial. Trials generate 1-3 best implicit ventriculum contraction model network solutions. The process is repeated to accumulate distinct best network solutions (best model scenarios) per DAG inspired model.

**Figure 2.**
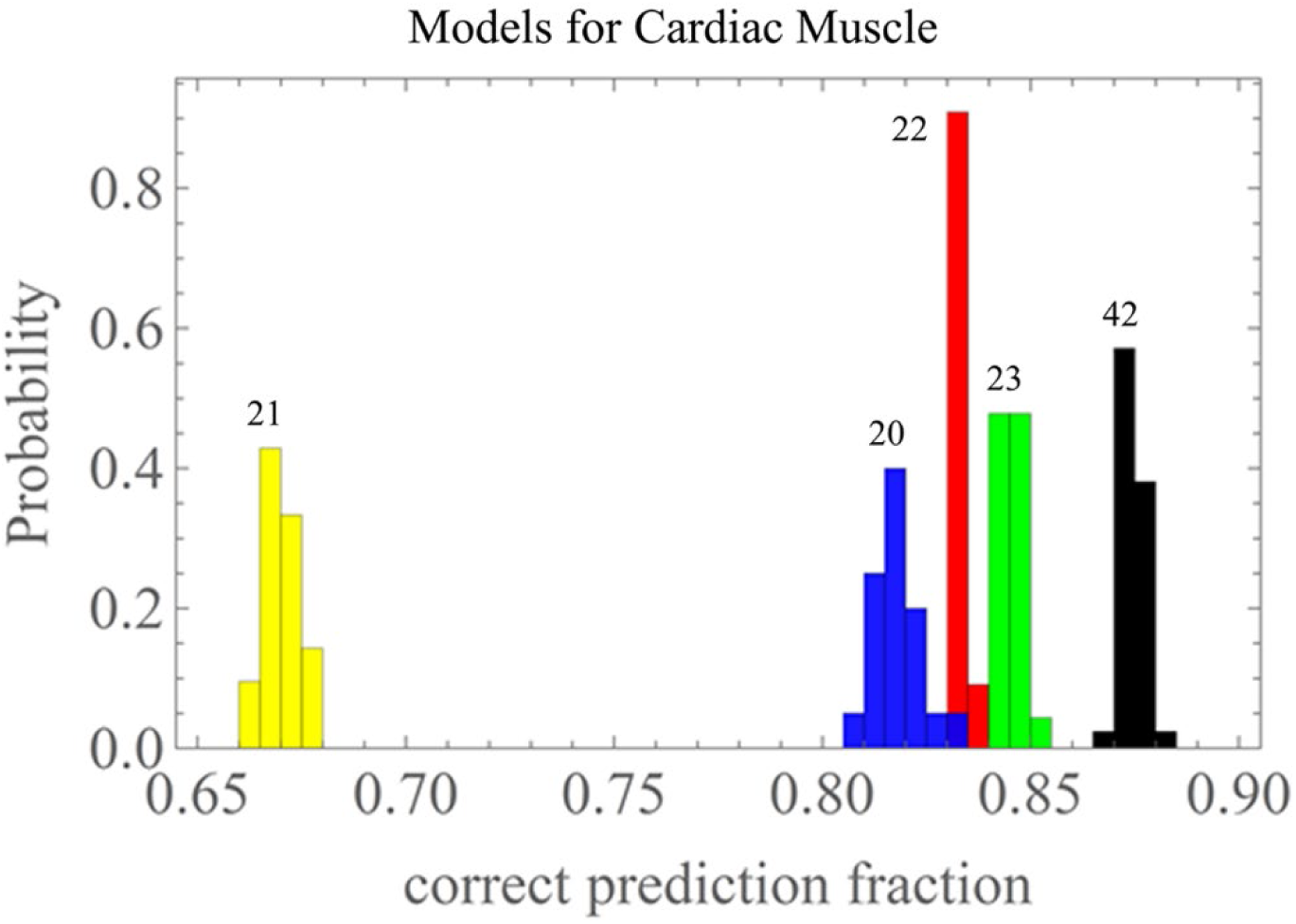
Correct prediction fractions (CPFs) for optimized DAG inspired general contraction Models: 6 (black), 7 (red), 8 (green), 9 (blue), 10 (yellow). CPF=1 is perfect model equivalence to validation data. Total probability for each model is normalized to 1. Numbers atop the probability histograms are scenarios count. Model 6 performs best with CPF >0.87.

CPFs in **Figure 2** are the best model scenarios grouped by DAG design. The histograms indicate a hard upper limit on correct predictions that cleanly identifies Model 6 (black) as best. It had the >0.88 CPF twice in the 42 scenarios and correct fractions >0.87 predominate. Model 6 scenarios correctly identified >87% of the phenotype and pathogenicity outputs in the fulfilled 6ddps. Model 10 is the poorest performer. Models 7-9 form a cluster near Model 6 but with CPF ≤0.85. The best Model 6 scenarios included hidden layer depths of 1-2 (for both N1 and N2) and widths up to 9 x (input nodes). Model 6 is the best choice for identifying co-domains in complexed βmys/MYBPC3. Model 6 scenarios outnumber those from the other models in **Figure 2** because their number was expanded for good statistics in the application described next after it was determined they were the best scenarios.

### Complex βmys/MYBPC3

**Figure 3** indicates homology (βmys) and linearized models (βmys and MYBPC3), many of their structural/functional domain locations, and the two letter abbreviations for the domains involved in co-domain interactions described below. Supplementary Information (SI) **Figure S1** lists all domains defined for the βmys/MYBPC3 complex.

**Figure 3.**
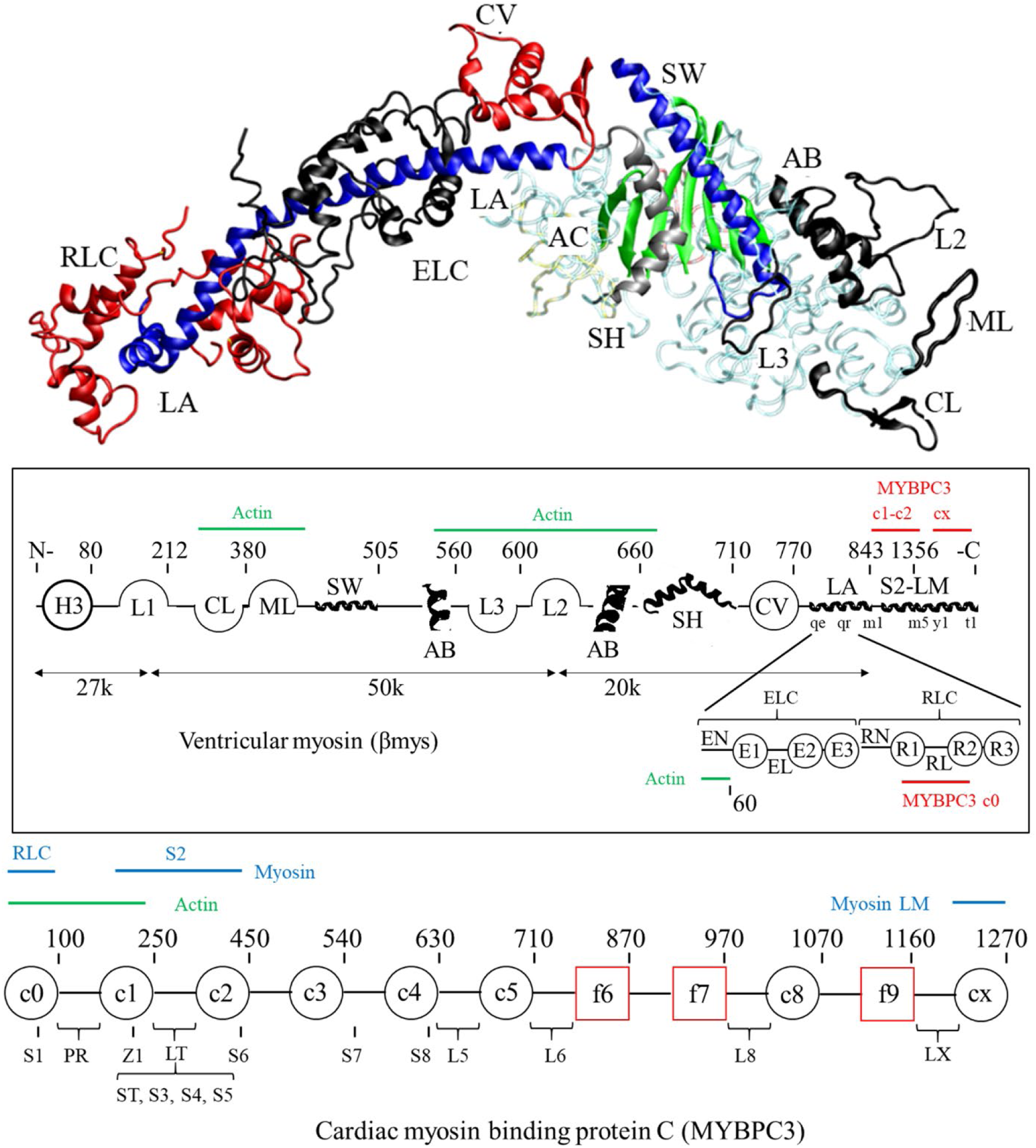
Homology and linearized models for the βmys structure (top and middle) and the linearized model representing MYBPC3 (bottom), identifying most domains defined in SI **Figure S1**. The linearized myosin diagram (middle diagram inside box) does not indicate the active site (ac), OM binding site (om), and mesa (me) because they occupy multiple regions in the linearized representation. Myosin light chains appear below the heavy chain. Several known MYBPC3 and actin binding sites are indicated in red and green above the heavy chain and below the light chains. The MYBPC3 diagram (bottom) has 8 Ig-like domains (black circles) and 3 fibronectin-like domains (red squares). Serine phosphorylation sites: S1 (Ser47), ST (Ser273), S3 (Ser282), S4 (Ser302), and S5-S7 and a threonine phosphorylation site (S8) are indicated below the chain. Domain linkers of interest include the proline rich linker (PR) and two (LT) containing a regulatory site. LT, L5, L6, and L8 are probable swivels. Z1 is a zinc binding site. Several known myosin RLC, S2 and LMM (blue) and actin binding (green) sites on MYBPC3 are indicated above the linearized model.

**Figure 4** **panels a-c** show 2D-CG maps of combined synchronous and asynchronous cross-correlate amplitudes (co-domain amplitudes, see Methods eq. 9) identifying co-domains linking βmys with MYBPC3 in the βmys/MYBPC3 complex for FHC, DCM, and LVN phenotypes. Maps are constructed as described in Methods Section *2D correlation testing and significance*. Each pixel in a map corresponds to a single co-domain pair. All DAG model designs were involved in optimization trials, however as expected from the CPFs in **Figure 2**, best model scenarios involved the Model 6 design exclusively.

**Figure 4.**
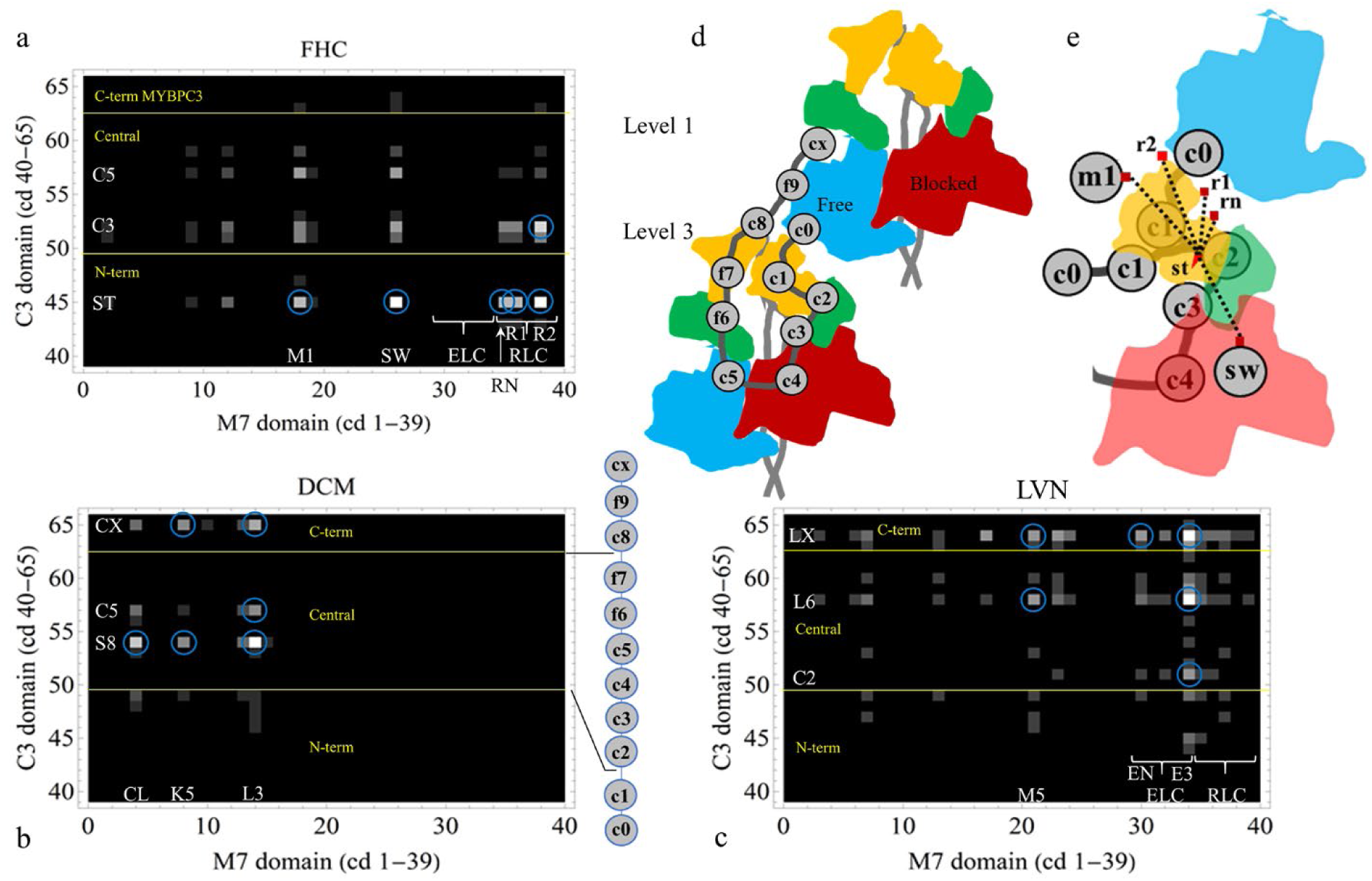
Cross-correlated SNV probability maps linking co-domains spanning the βmys/MYBPC3 complex for FHC, DCM, and LVN phenotypes (**a-c**), hypothetical model for two myosin dimers and MYBPC3 on the surface of the thick filament in relaxed ventriculum (**d**), and (**e**) selected portions of structure in (**d**) after FHC-causing SNV binds to MYBPC2 at Serine 2 domain (ST) in linker 2 (LT). (**a**) Cross-correlate intensity indicated by the grey scale intensity against 0 (black) for FHC. Axes have βmys (M7) domains (1-39 along abscissa) and MYBPC3 (C3) domains (40-65 along ordinate). Selected co-domains are labeled with their two letter codes. Domain indexing and two letter codes from SI **Figure S1**. Yellow horizontal lines and legends refer to MYBPC3 segments: N-terminus (C0-C1), Central segment (C2-F7) and C-terminus (C8-CX) also shown by the simplified MYBPC3 model beside the 2D map in panel **b**. Blue circles indicate the 6 most significant co-domains. (**b-c**) Notation identical to panel **a** but for DCM and LVN phenotypes, respectively. (**d**) Cartoon representation of two βmys dimers corresponding to crown Levels 1 and 3 along a quasihelical strand on the thick filament from EM reconstruction [1] and MYBPC3 after Ponnam and Kampourakis [2] with modifications. Myosin heads are in blocked (red) and free (blue) conformations with light chains ELC (green) and RLC (orange). Myosin S2 shown in lighter grey scale. MYBPC3 domains are numbered as in Figure 3 and linkers shown it darker gray scale. (**e**) model structure in (**d**) in the presence FHC-causing SNV bound to MYBPC3 ST (red triangle) showing hypothetical MYBPC3 N-terminus displacement by the SNV at ST and virtual co-domains linking ST and βmys RLC, switch 2 helix (SW), and MYBPC3 binding site on myosin S2 (M1). Red squares indicate FHC-causing SNV positions on βmys and dashed black lines the virtual co-domain links.

Co-domain probability amplitudes are distributed normally about zero as described in Methods. Outliers include the 6 most outstanding indicated by the blue circles in the maps. They fall >78, >66, and >14 standard deviations from their mean value for the FHC, DCM, and LVN phenotypes, respectively. Co-domain outliers identify the most certain pairs coordinating interactions between βmys and MYBPC3. Co-domain footprints linking collectively motor domain and lever-arm bound light chains in βmys with the N-terminal segment (C0-C1), central segment (C2-F7), and C-terminal segment (C8-CX) domains in MYBPC3 clearly differentiate the three phenotypes.

Sub-phenotypes {hx, h1, h4, h8}, {ds, d1, dc}, and {lc, lv, lw} (SI **Figure S2**) comprise the broader FHC, DCM, and LVN phenotype categories, respectively. All sub-phenotypes denote exclusively SNV modification in βmys (MYH7, MYL2, MYL3) or MYBPC3. Common FHC, DCM, and LVN {population, domain} pairs (Methods eqs. 3-5) were selected exclusively for analysis to rationalize their comparison. They are comprised from 1562±83, 514±38, and 1624±89 different SNVs for a total of 3706±85 (mean ± standard deviation) for N=42 model scenarios. SNV counts vary over the 42 unique contraction models as their prediction for unknown phenotypes differ. SNV coverage per domain+linker (C-terminal segment has 1 less linker that the others) is heaviest for the N-terminal segment containing 958±44, compared to central and C-terminal segments containing 1890±50 and 858±28. SNV coverage is extensive for the three phenotype categories.

Co-domain based SNVs in the βmys/MYBPC3 complex (or in any complex) can detect co-domains (eg., bright pixel above threshold inside blue circles in **Figure 4a-c**), or, detect and disrupt co-domains. The latter holds because detection does not require an intact co-domain. A co-domain is detected because of the larger product of probabilities for SNVs occupying either site in the two co-domain members (Methods eqs. 3-5). Single residue variants probably alter a co-domain’s ability to communicate inferences between proteins rather than disrupt their bi-molecular association but the latter is not excluded.

The cartoon model in **Figure 4** **panel d** has the βmys/MYBPC3 complex, after Ponnam and Kampourakis (see Figure 5 in [2]), qualitatively summarizing findings from 2D-CG. Results in **Figure 4d** differ from the earlier work. βmys dimers form the thick filament in human ventricular muscle. βmys heads have motor domain, lever arm, and light chains ELC (green) and RLC (orange) with heads lying on the filament surface. The top and bottom βmys dimers correspond to crown Levels 1 and 3 along a quasihelical strand on the thick filament as described by AL-Khayat et al. from EM reconstruction [1]. Motor domains within each dimer form a complex of free (blue) and blocked (red) heads. The βmys dimers in **Figure 4d** link to each other at a contact from the Level 1 dimer free motor domain (blue) to the Level 3 dimer RLC on the blocked head (red) to form the SRX structure.

FHC linked co-domains involve βmys RLC, switch 2 helix (SW), and a MYBPC3 binding site on myosin S2 (M1) with the MYBPC3 Serine 2 (ST) in the linker 2 (LT) regulatory domain (**Figure 3**). Another co-domain link has βmys RLC subdomain R2 with MYBPC3 C3. Involved MYBPC3 domains are from the N-terminus and central segments implying, with inspection of **Figure 4d**, that FHC linked co-domains detect interactions between MYBPC3 and the free myosin head at the Level 3 myosin dimer along the quasihelical strand. Level 1 dimer interactions are not detected until <21 standard deviations from the mean value for FHC. Co-domain interactions between βmys and MYBPC3 for the Level 1 myosin dimer in the FHC affected muscle are exceedingly rare (disrupted) compared to those engaging the Level 3 dimer. It appears that FHC linked SNVs disrupt coordination between the Level 1 myosin dimer and the MYBPC2 C-terminus but that SRX disruptive co-domain SNVs are not in the MYBPC3 C-terminus (because nothing is detected there for the FHC affected complex).

DCM linked co-domains coordinate βmys actin binding sites including C-loop (CL) [17–19], Loop 3 (L3), and the 50kDa proteolytic fragment (K5) of βmys in the motor domain with the central segment of MYBPC3 including Serine 8 (S8) and C5, and, with the C-terminus segment at CX. S8 and C5 are at opposite ends of linker 5 (L5) in the central segment. The (βmys-MYBPC3) CL-CX co-domain appears in the map but falls just short of the significance threshold (as indicated by the blue circles). Co-domains containing the MYBPC3 central segment domains imply interaction with the blocked and free heads of the Level 3 myosin dimer. Co-domains involving the C-terminal CX implies interaction between MYBPC3 and the free head of the Level 1 myosin dimer and as indicated by the EM reconstruction [1]. DCM linked co-domains imply MYBPC3 interacts with both myosin dimers. Data from DCM SNVs do not (by themselves) rule out the possibility that DCM SNVs detected in the MYPC3 C-terminus disturb its interaction with the Level 1 myosin dimer. However, when combined with FHC SNV data showing SRX disruptive co-domain SNVs are not in the MYBPC3 C-terminus and that these findings are based on 2D-CG analysis of common FHC, DCM, and LVN {population, domain} pairs, the evidence favors a conclusion that DCM SNVs do not disturb the MYPC3 C-terminus/Level 1 myosin dimer interaction.

LVN linked co-domains target βmys ELC and M5 (a MYBPC3 binding site on LMM) principally with linker 10 (LX) in the MYBPC3 C-terminus segment, and, the βmys ELC with MYBPC3 C2 and linker 6 (L6) in the central section. Co-domains containing the MYBPC3 C-terminus domains suggests interaction between MYBPC3 and the free myosin head of the Level 1 myosin dimer. The other co-domains in central section of MYBPC3 implies interaction between MYBPC3 and both free and blocked heads of the Level 3 myosin dimer. LVN linked co-domains suggest MYBPC3 interacts with both myosin dimers. The argument supporting the idea that LNV SNVs do not disturb the MYPC3 C-terminus/Level 1 myosin dimer interaction parallels that given in the previous paragraph for DCM SNVs.

DCM and LVN co-domain footprints have interactions linking both the C-terminal segment (LX and CX) and central segment (S8-c5 and L6) with the myosin heads implying interactions with both Level 1 and Level 3 myosin dimers along a quasihelical strand in the thick filament forming the SRX structure. The co-domain footprint for FHC does not have a detectable Level 1 dimer interaction thus breaking SRX structure. Every co-domain linkage in the βmys/MYBPC3 complex is detectable by the SNVs in the database using 2D-CG but not all of them can be represented in the **Figure 4d** model for SRX myosin. For example, thick filament structures other than SRX are present in the human populations represented in the database used for this study. Mindful of these known-unknowns, co-domain linkages in the βmys/MYBPC3 complex are qualitatively represented by their position in the **Figure 4d** model.

## DISCUSSION

Thick filament structure surmised from human ventricular muscle under relaxing conditions [1] has myosin heads apparently in the off-state [20] and super-relaxed because they associate with the thick filament unable to interact with actin [21]. SRX has myosin dimers stabilized on the thick filament along the quasihelical strand where inter-dimer contacts form. The disordered relaxed state (DRX) has myosin heads moving away from the thick filament to a position where they can contact actin [22]. DRX myosin dimers do not form stable inter-dimer contacts as they are mobile. In DRX, MYBPC3 remains tethered to the thick filament at CX but loses contact with the Level 1 myosin dimer in favor of the Level 3 dimer in **Figure 4d** since otherwise Levels 1 and 3 myosin dimers would be unable to move away from the thick filament. DCM and LVN-causing SNVs identify co-domain linkages spanning different myosin dimers across two levels in the quasihelical strand suggesting these phenotypes do not disturb a dominant SRX structure in a functioning human heart. In contrast, FHC-causing SNVs identify co-domain linkages in just Level 3 dimers on the quasihelical strand suggesting they disturb SRX in agreement with earlier work associating FHC with the SRX disruption in humans and in animal models [21, 23, 24].

SRX destabilization by FHC-causing SNVs does not involve co-domains between βmys and the MYBPC3 C-terminus because these interactions would be detected in the 2D-CG map in **Figure 4a** (recall that SNV induced co-domain destabilization does not prevent their detection). It supports the notion that co-domains between βmys and the MYBPC3 C-terminus detected by DCM and LVN-causing SNVs detect but do not destabilize SRX. SRX disruption must involve a βmys domain hence SNVs in MYBPC3 that cause FHC must do so by destabilizing SRX through an indirect (direct would be via the MYBPC3 C-terminus) pathway such as an allosteric mechanism. A suitable allosteric mechanism was described previously as a virtual-mechanism co-domain that would also be detected by 2D-CG (see Methods section *Co-domain interaction by real or virtual mechanisms* or [10]). Candidate virtual co-domains would include those for which the MYBPC3 domain interacts with multiple βmys domains.

FHC-causing SNVs in the MYBPC3 N-terminal segment at Serine 2 (ST) domain forms 5 of the 6 most significant co-domain associations with βmys (**Figure 4a**). Three with sub-domains of RLC (RN, R1, and R2) and two with βmys sites in the lever arm adjacent to RLC (M1) and the motor domain (SW). These potential virtual co-domain links are indicated in **Figure 4e** by dashed black lines. The SNV at ST (red triangle) induces a proposed conformation change in the linker 2 (LT) swivel [3] disturbing the N-terminus and revealing virtual βmys co-domain sites on RLC, M1, and SW. The scale of the MYBPC3 N-terminus segment displacement by the SNV at ST is unknown. Displacement shown in **Figure 4e** is for illustration.

In the absence of the SNV at ST, real co-domains between unidentified sites on MYBPC3 in the N-terminus segment and βmys domains {RN, R1, R2, M1, SW} are proposed to stabilize the Level 1-Level 3 myosin dimer interaction at the free head (Level 1)-RLC (Level 3) interface favoring SRX. The LVN 2D-CG map (**Figure 4c**) has below significance threshold co-domains appearing in the MYBPC3 N-terminus segment corresponding to interactions between MYBPC3 (ST, LT) and βmys RLC (RN). These might identify two of the unidentified MYBPC3 members of the real co-domains stabilizing SRX with their counterparts in βmys. In this general contraction model (Model 6), ST and sites closer to the N-terminus from ST but within the regulatory domain LT on the MYBPC3 side form co-domains with βmys domains {RN, R1, R2, M1, SW} in the Level 3 myosin dimer. It appears then that MYBPC3 ST forms both real and virtual co-domains with βmys. Only real co-domains correspond to a physical contact. Co-domain RN on the βmys side contains the RLC phosphorylation site at Serine 15. MYBPC3 N-terminus regulates contraction [25] as does its βmys RLC co-domain. RLC phosphorylation affects SRX stability [26, 27] and βmys step-size [28] in the native system. Other links between RLC and MYBPC3 are already established [29].

In the presence of the SNV at ST, real co-domains at ST and sites closer to the N-terminus from ST but within the linker 2 (LT) regulatory domain on the MYBPC3 side, are disturbed. It destabilizes the Level 1-Level 3 myosin dimer interaction at the free head (Level 1)-RLC (Level 3) interface. The destabilization potentially mimics the native mechanism for disrupting SRX by RN phosphorylation. RLC has several variants implicated in FHC disease [30–32] including one in R1 that stabilizes SRX in mice [33]. The latter implies there are other molecular mechanisms for inheritable FHC.

The ST-RLC co-domain interactions (characteristic to a destabilized SRX in FHC) are not detected at the highest significance level in DCM or LVN species again implying their C-terminus SNVs are SRX detectors but not disruptors such that the MYBPC3 C-terminus segment SNVs do not influence SRX stability. The DCM result agrees with a previous analysis of myosin variants [34, 35]. The 2D-CG map for FHC SNVs appears to eliminate the free head (Level 1)-RLC (Level 3) contact destabilizing the SRX conformation. Other work in animal models suggests FHC SNVs do not eliminate SRX but rather down modulates its presence [24]. The latter could reflect incomplete allele replacement in SNV affected models while the loss of the free head (Level 1)-RLC (Level 3) contact seen in the human genetic data does so because detection is specific to SNV modified βmys/MYBPC3 complexes.

Co-domains from each phenotype engage the MYBPC3 central hinge in and around C5 [3]. It is positioned to influence the relationship between free and blocked myosin heads (**Figure 4d**), perhaps by subtracting rigidity from their association, when the βmys/MYBPC3 complex leaves the thick filament because of RLC phosphorylation [26]. The 2D-CG results for DCM and LVN implicated SNVs indicate that the most significant co-domains interactions avoid the N-terminus section of MYPBC3 drawing a sharp contrast with the FHC implicated SNVs. DCM and LVN similarly retain their SRX filament structure while having different SNV induced impairments targeting βmys actin binding (DCM) and mainly ELC (LVN). Different targets imply different mechanisms for disease. Acto-myosin interaction impairment will affect the ability of the muscle to produce force while ELC modification will likely have a more nuanced impact. The LVN phenotype suggests sarcomere assembly in the muscle is affected.

Another objective undertaken was exploring how to interpret global neural networks models for contraction mechanism (those in **Figure 1**) with basic natural language characteristics. Models in **Figure 1** separate data (represented in circles) into inputs and outputs. Inputs {cd,re,po,af} are basic and reliable SNV characteristics giving rise to outputs {ph,pa} that are more difficult to ascertain, sometimes inconsistent, and reported less frequently. Influences (represented in arrows) assign pathways that are permuted in the models to measure their ability to mimic the system. The models evaluated in **Figure 2** show that phenotype input to the pathogenicity net chain at N2, in Model 6, is essential to model accuracy and superior to the other configurations.

All models have the medically relevant natural language characteristics phenotype and pathogenicity explicitly embedded in the DAG. Model 6 is the starting point to explore emergent basic structural/functional characteristics implicit in it.

Deeper probing into Model 6 characteristics tested hidden layer depth and width in network chains (N1 and N2, **Figure 1**) to find best model scenarios. The latter rarely had hidden layer depths different from 1 and never went above 3 in either of the network chains. This was unexpected. On the other hand, width was more varied for both N1 and N2, the network chains managing phenotype and pathogenicity. Width, expressed as multiples of input nodes to the network chain, ranged from 1-8 for N1 and 1-10 for N2. Width adjustment was likely locating an optimum rank set of input characteristics encoded by the four input variables {cd,re,po,af} also including *ph* for N2. It is larger than the sum of their inputs because, for instance allele frequency (af), depends on other inputs like population (po). It implies inputs are not independent because cardiac muscle physiology integrates βmys/MYBPC3 functionality with the wider system involving genetic background [9]. These deeper dependencies are coded in hidden layer width and depth that, when deciphered, emerge as a basic natural language mechanism characteristic.

Biochemical and mechanical characterization of the contractile system would likewise help with recognizing natural language contraction mechanism characteristics from a general neural net model, however, that is a costly venture and unlikely to be pursued on a world-wide scale unlike genetic screening. Instead, the neural network is a shortcut supplying unknown phenotypes and pathogenicities interpreted, together with the known instances, using Bayes network probabilities implied by the DAG. They are the means to decipher population dependencies in the genetic screening data to map influences mediated by SNV modified co-domains using 2D-CG [10]. In this application 2D-CG informs the thick filament structural state for FHC, DCM, and LVN phenotypes as destabilized-SRX, SRX, and SRX, respectively, implying a new output defined as thick filament structural state (fs) enlarging 6ddps to 7ddps as *fs* joins *ph* and *pa*. Filament state quantitation discovered with the focus on FHC, DCM, and LVN will eventually encompass the other phenotypes in the database. Filament state is the emergent structurally relevant natural language characteristic incorporated into the “Model 11” DAG prototype in development.

In the bigger picture, co-domains define inter-protein influence pathways that recall energy transduction pathways in the myosin ATPase mechanism driving tension generation [36]. Mapping influences mediated by SNV modified *intra*-myosin domains identifiable using 2D-CG would involve discoverable transduction pathways (tp) augmenting the existing large biochemical datasets. Transduction pathway could be another emergent structurally relevant natural language characteristic on the horizon to be incorporated into future general network contraction models.

## CONCLUSION

Human ventriculum βmys powers contraction sometimes while complexed with MYBPC3. The latter regulates βmys activity through inter-protein contacts. SNVs change protein sequences in βmys or MYBPC3 and cause inheritable heart disease. When a SNV modified domain locates to an inter-protein contact it affects complex coordination. Domains involved, one in βmys and the other in MYBPC3, form coordinated domains (co-domains). Co-domains are bilateral implying the potential for a shared impact from SNV modification in either domain suggesting their joint response to a common perturbation assigns location. Human population genetic divergence is the common systemic perturbation. A neural/Bayes network design general contraction model reveals SNV probabilities for SNV characteristics and specifies correlations between domain members revealing co-domain locations in three common human heart diseases caused by SNVs, FHC, DCM, and LVN. Co-domain maps for DCM and LVN link MYBPC3 central (C2-F7) and C-terminal (C8-CX) segments with myosin heads implying interactions with both Level 1 and Level 3 dimers along a quasihelical strand in the thick filament. These myosin dimers form SRX. The FHC co-domain map does not involve the Level 1 dimer implying these myosins do not form the SRX. Comparing co-domain maps for the FHC, DCM, and LVN phenotypes suggests the SRX disruption mechanism involves a co-domain between the MYBPC3 regulatory and RLC N-terminus domains. Best general contraction model scenarios found for the FHC, DCM, and LVN systems were explored with the purpose to understand how to interpret them with basic natural language characteristics. They emerge from dependencies among inputs coded in hidden layer width and depth when they are deciphered using 2D-CG. In this application 2D-CG informs the thick filament structural state for FHC, DCM, and LVN phenotypes defining the thick filament structural state output (fs) joining phenotype and pathogenicity.

## METHODS

### SNV data retrieval

SI **Figure S3** outlines the protocol for SNV data retrieval from the National Center for Biotechnology Information (NCBI). SNV reference number (rs#) identifies the affected gene position and one or more alleles corresponding to synonymous and missense variants. Missense variants are selected providing 4 input parameters to the models (**Figure 1**) defining the protein domain affected (cd), residue substitution (re), population group (po), and allele frequency (af) for the SNV. Associated clinical data provides 2 output parameters phenotype (ph) and pathogenicity (pa). Input parameters are always known quantities. Output parameters are frequently unknown or contradictory among the data submitters. Input and output parameters form a 6-dimensional data point (6ddp). The 6ddps are combined into a matrix collecting like data from each SNV in the record. The subset of 6ddp’s containing unknown output parameters (ph, pa, or both) is the unfulfilled 6ddp matrix. The subset of 6ddp’s containing all known parameters form the fulfilled 6ddp matrix.

### Neural/Bayes network configuration

Directed acyclic graphs (DAGs) in **Figure 1** show the general neural network models for contraction tested for associating the inputs with outputs as described [10]. **Figure 1** Model 6 was used previously [8, 10, 37]. It has *ph* causal for *pa*. Model 8 reverses roles of *ph* and *pa* while Model 7 eliminates causality between output variables *ph* and *pa*. Models 9 isolates causality from input parameters to *ph* then from *ph* to *pa*. Model 10 reverses the roles for *ph* and *pa* from Model 9. These trial models represent different versions of muscle and muscle disease mechanisms that are tested for suitability by comparing their accuracy for representing SNV data as described in the section, Neural Network Validation.

SI **Figure S1** indicates the protein domains (cd), 2 letter abbreviations, and sequence location. A protein complex made from the four genes, MYH7, MYL2, MYL3, and MYBPC3 has domains from 65 functional sites as described [10]. Every SNV in the database has an assigned domain. **Figure 3** shows linear representations of βmys and MYBPC3 indicating some mutual binding sites and the locations of most domains listed in SI **Figure S1**. In the future, finer grained functional domain assignment in the complex will enhance spatial accuracy for co-domain assignments.

Residue substitution (re) refers to the nonsynonymous SNV reference and substituted residue (ref/sub) pairs. Residue abbreviations are the standard 1 letter code. Ref/sub combinations have 420 possibilities for 21 amino acids. Nonsynonymous SNV substitutions for the βmys/MYBPC3 complex in the NCBI database has 147 unique ref/sub pairs. It is a large expansion in neural network complexity compared to the earlier work achieving substantially higher accuracy SNV outcome prediction than previously [10].

Human population group and allele frequency fill out the independent parameters in the network. SI **Figure S4** indicates populations and their 3 letter abbreviations. Allele frequency (af) is a continuous variable in the database on the interval 0 ≤ af ≤ 1 for 1 meaning all alleles are substituted by the SNV. These data are divided into two discrete categories of ≤1% (category 0) or >1% (category 1) for this application.

The NCBI SNP database has 17 phenotype data classifications for cardiovascular disease pertaining to βmys and MYBPC3 variants. SI **Figure S2** lists names and two letter codes. Some phenotypes associate with both βmys and MYBPC3 SNVs. Data submissions from different providers occasionally conflict for a given SNV. They are assigned from the pool when there is a clear consensus. In all other cases the unknown category is assigned.

Pathogenicity data classifications include pathogenic (pt), likely pathogenic (lp), benign (be), likely benign (lb), and unknown (uk). Data submissions from different providers occasionally conflict for a given SNV. They are assigned from the pool when there is a clear consensus. In all other cases the unknown category is assigned.

### Neural network training and validation

Neural networks in **Figure 1** model structure/function influences from disease and imitate the DAG pathways linking inputs (cd,re,po,af) to outputs (ph,pa) [8]. Prototype neural networks for Models 6 and 8 are depicted in the figure. They are tasked with predicting unknown phenotype and pathogenicity from the input parameters defining a SNV.

The top model optimization layer (already described in Results) implies searching through 1-6 hidden layer depths, and the (1-10) x (input node count) hidden layer widths for the 5 DAG inspired model architectures (**Figure 1**). Each model optimization run is called a trial. Trials generate 1-3 best implicit ventriculum contraction model network solutions called general contraction model scenarios. The process is repeated to accumulate several distinct best scenarios. A trial consists of the steps outlined as follows.

Underneath the top layer is the second optimization layer concerned with the neural networks in each model with specified DAG inspired model architecture and hidden layer dimensions. These specific models are trained and validated using the validating dataset composed of all the fulfilled 6ddps. The training dataset is half of the validating data with members chosen randomly but subject to the constraint that each output is represented in the training dataset except when their representation in the validating dataset is <2 occurrences. Learnable parameters in the networks are randomly initialized. Weight initializations are normally distributed with zero mean and standard deviation of (1/n)^½^ (for n inputs). Bias is initialized to zero. Loss, the number of incorrect predictions from inputs for outputs, is minimized by adjusting learnable parameters in the neural networks using stochastic gradient descent. The validation dataset tests each trained network using a preconfigured loss layer from Mathematica (CrossEntropyLossLayer) appropriate for output from the softmax layer (**Figure 1**). Inclusion of the preconfigured dropout layer from Mathematics routines (**Figure 1**) mitigates overfitting. Overfitting is likewise avoided by terminating training when the validating dataset loss begins to increase following the initial descent during iterative loss minimization. The latter overfitting control is adjusted as appropriate for the model form (in Models 6-10), hidden layer depth and width, i.e., model complexity. The full second optimization layer consists of several steps with stochastic initial condition choices that are sampled 4-10 times with the different initial conditions generated by their stochasticity. The ability to correctly classify SNVs measures the overall suitability of each model scenario as shown by the CPF (**Figure 2**).

### Bayes network modeling of complexed βmys/MYBPC3 transduction mechanism

**Figure 1** shows DAGs for the Neural/Bayes network model. Arrows imply a direction for influence hence the domain, residue substitution, population, and allele frequency assignment collectively imply probability for phenotype and pathogenicity. Supplementary Information (SI) contains fulfilled and unfulfilled 6ddps (6ddpdatasetAll.xls with 26,056 variants) and fulfilled only 6ddps (6ddpdatasetFull.xls with 4,093 variants) for complexed βmys/MYBPC3. The full dataset, with unfulfilled entries fulfilled in turn by each of the best model scenarios (42 in this case), define conditional probabilities for the systems in the form of conditional probability tables (CPTs). The product of conditional probabilities on the right defines the joint probability density on the left in,

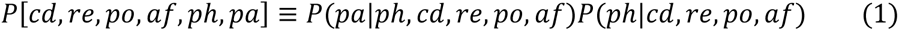

Calculating SNV probability for domain *i* in population *j* with phenotype *k* uses joint probability densities,

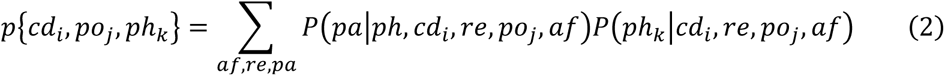

where summation is over all values for allele frequency, residue substitution, and pathogenicity. Summing over phenotypes defining FHC (hx, h1, h4, and h8), DCM (ds, d1, dc), or LVN (lc, lv, and lw) specifies SNV probability representing each ailment in eqs. 3-5.

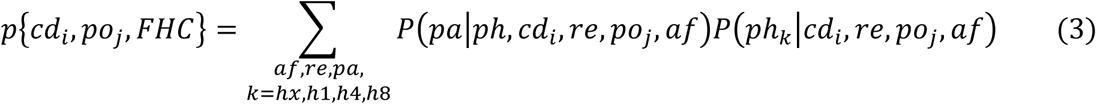

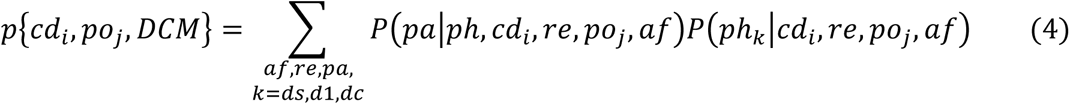

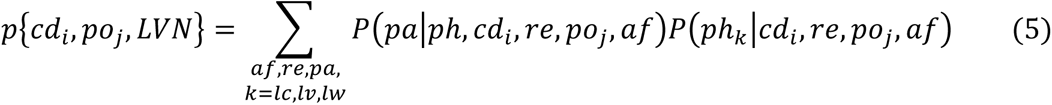

### 2D correlation testing and significance

SNVs from human population *po_j_* reside in a protein domain *cd_i_* with probability given by eq. 3, 4, or 5 depending on disease. Eqs. 3-5 expressions populate the synchronous and asynchronous generalized 2D correlation intensities [14] and as described [10],

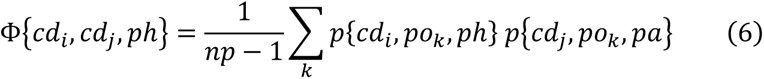

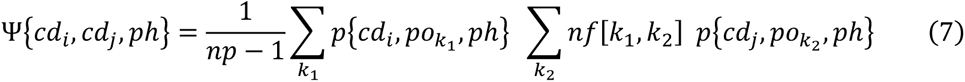

for,

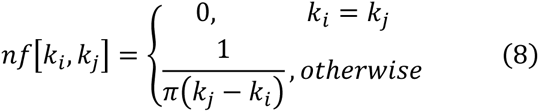

Φ in eq. 6 gives synchronous, and Ψ in eq. 7 asynchronous, 2D correlation intensities for *np* the number of populations represented by *po* and phenotypes ph = FHC, DCM, or LVN. The 2D synchronous intensity map will identify domain pairs whose SNV probabilities correlate or anti-correlate for identical populations while the 2D asynchronous intensity map identifies the leading and lagging co-domain member over populations ordered by their genetic divergence.

The most significant combined synchronous and asynchronous co-domain interaction cross-correlates are the largest elements in the array,

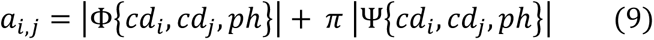

for Φ and Ψ from eqs. 6 and 7, *cd_i_* and *cd_j_* the SNV containing functional domains in βmys and MYBPC3, and *ph* their common phenotype (FHC, DCM, or LVN). Array element *a_i,j_* is the co-domain cross-correlate amplitude given by the product of probabilities for SNVs in each domain member of the co-domain weighted by synchronous or asynchronous coupling to human populations. Quantity π multiplying the asynchronous cross-correlates (Ψ) in eq. 9 balances weighting for synchronous (Φ) with nearest neighbor population asynchronous cross-correlates. Elements *a_i,_*_j_ are ≥ 0. They are combined with *−a_i,j_* and their distribution visualized in a histogram. A normal distribution approximates the result. Absolute distance from the mean expressed in multiples of standard deviation indicates a measure of co-domain cross-correlate amplitude significance. Co-domain cross-correlate amplitude significance increases with distance from amplitude mean.

Data obtained from calculation of protein domain SNV probability correlations is represented in 2D plots showing the amplitudes in eq. 9 for βmys (abbreviated M7) on the abscissa and MYBPC3 (C3) domains on the ordinate, in the order given in SI **Figure S1**. Domains are discrete entities hence the 2D plots resemble a pixelated image with grayscale representing intensity. Each pixel in the maps corresponds to a co-domain pair. Only inter-protein (co-domain) correlations are indicated.

### Population genetic divergence proxy

Worldwide human genetic divergence is attributed to migration from a single origin in East Africa based on the serial founder effect [11–13]. The serial founder effect explains the observed linear divergence decrease with human migration distance over the earth’s surface. Migration distance is therefore a good proxy for genetic divergence variation.

Estimates for migration distances on earth’s surface are calculated exactly as described [10]. Distance estimates (and genetic divergence variation) from the fitted line for populations in SI **Figure S5** are equally spaced over the Population Index parameter as needed for the co-domain 2D-CG formalism.

### Co-domain interaction by real or virtual mechanisms

A co-domain pair from two proteins, A and B, in complex implies two functional domains (one in A and one in B) where SNV modification probabilities cross-correlate over human populations. It occurs by two mechanisms named real and virtual [10]. The real mechanism is indicated for the βmys/MYBPC3 complex in SI **Figure S6 panel a** by the solid green lines linking two co-domain pairs. They represent a direct through-space (real) interaction within the βmys/MYBPC3 complex and requires two SNVs for detection. In this case SNV_1_ and SNV _2_ for S3 and CL, or, SNV_3_ and SNV _4_ for C5 and ML. The virtual mechanism, indicated in SI **Figure S6 panel b** with the dashed green lines, implies the SNV at L8 (SNV_5_) disturbs the bend in the MYBPC3 at this site that disrupts real S3-CL and C5-ML co-domains. The virtual mechanism has a SNV altered functional domain that perturbs a real co-domain contact. The real co-domain contact does not involve L8. Real co-domains involve just a pair of domains while virtual co-domains could involve multiple domain interactions. If multiple interactions are detected, for example at L8-CL and L8-ML the virtual mechanism is likely.

## Supporting information

All SNVs

Fulfilled SNVs

## Acknowledgment

The author thanks Katalin Ajtai for scientific discussion and critical review of the manuscript.

## Supplementary Information

Supplementary information consists of a document with Figures S1-S6 and two data sets for complex βmys/MYPBC3. Data sets in files 6ddpdatasetAll.xls and 6ddpdatasetFull.xls contain all and fulfilled only 6ddps.

## Declarations

### Data Availability

All data generated or analyzed during this study are included in this published article and its supplementary information files.

### Code availability

All computer coding is in Mathematica and available from the author.

### Author contribution

TPB is the sole author and responsible for the content of the paper. The author read and approved the final manuscript.

