## Supplementary Information for "Variants identify sarcomere inter-protein contacts distinguishing inheritable cardiac muscle diseases"

Thomas P. Burghardt

Department of Biochemistry and Molecular Biology

200 First St. SW

Mayo Clinic Rochester

Rochester, MN 55905

https://orcid.org/0000-0003-3119-4074

| domain (cd) | seq | index |
|---|---|---|
| M7 actin binding (ab) | <525-563, 647-655> | 1 |
| M7 active site (ac) | <114-125, 167-178, 244-253, 260-267, 453-465, 666-673> | 2 |
| M7 Blocked Head/Converter (bh) | various 295-397 | 3 |
| M7 C-loop (cl) | 359-377 | 4 |
| M7 Converter (cv) | 711-768 | 5 |
| M7 SH3 (h3) | 28-78 | 6 |
| M7 20k (k2) | 634-843 | 7 |
| M7 50k (k5) | 209-633 | 8 |
| M7 27k (k7) | Nterm-208 | 9 |
| M7 Lever-arm (la) | 769-843 | 10 |
| M7 LMM (lm) | 1357-Cterm | 11 |
| M7 Loop 1 (l1) | 202-212 | 12 |
| M7 Loop 2 (l2) | 622-646 | 13 |
| M7 Loop 3 (l3) | 564-576 | 14 |
| M7 MESA (me) | various 168-664 | 15 |
| M7 Myopathy-loop (ml) | 400-414 | 16 |
| M7 MESA-trail (mr) | various 172-917 | 17 |
| M7 S2-C1 mybpc3 (m1) | 844-857 | 18 |
| M7 S2-LT mybpc3 (m2) | 924-936 | 19 |
| M7 S2-C2 mybpc3 (m3) | 858-870 | 20 |
| M7 LMM-Cx (m5) | 1554-1581 | 21 |
| M7 OM binding (om) | various 91-712 | 22 |
| M7 IQ ELC (qe) | 788-798 | 23 |
| M7 IQ RLC (qr) | 814-824 | 24 |
| M7 SH1/SH2 hinge (sh) | 683-710 | 25 |
| M7 Switch 2 helix (sw) | 466-505 | 26 |
| M7 Subfragment 2 (s2) | 844-1356 | 27 |
| M7 LMM-titin (t1) | 1815-1831 | 28 |
| M7 LMM-myomesin (y1) | 1506-1674 | 29 |
| M7 N-term ELC (en) | Nterm-60 | 30 |
| M7 EF1 ELC (e1) | 49-86 | 31 |
| M7 linker ELC (el) | 87-127 | 32 |

| domain (cd) | seq | index |
|---|---|---|
| M7 EF2 ELC (e2) | 128-163 | 33 |
| M7 EF3 ELC (e3) | 164-Cterm | 34 |
| M7 N-term RLC (rn) | Nterm-23 | 35 |
| M7 EF1 RLC (r1) | 24-59 | 36 |
| M7 linker RLC (rl) | 60-94 | 37 |
| M7 EF2 RLC (r2) | 95-129 | 38 |
| M7 EF3 RLC (r3) | 130-166 | 39 |
| C3 phospho Ser 1 (s1) | 43-51 | 40 |
| C3 c0-Ig like (c0) | 1-101 | 41 |
| C3 proline rich (pr) | 102-152 | 42 |
| C3 zinc site 1 (z1) | <208, 210, 223, 225> | 43 |
| C3 c1-Ig like (c1) | 153-256 | 44 |
| C3 phospho Ser 2 (st) | 271-279 | 45 |
| C3 phospho Ser 3 (s3) | 280-288 | 46 |
| C3 phospho Ser 4 (s4) | 300-307 | 47 |
| C3 phospho Ser 5 (s5) | 308-315 | 48 |
| C3 linker c1-c2 (lt) | 257-361 | 49 |
| C3 phospho Ser 6 (s6) | 423-431 | 50 |
| C3 c2-Ig like (c2) | 362-452 | 51 |
| C3 c3-Ig like (c3) | 453-543 | 52 |
| C3 phospho Ser 7 (s7) | 546-554 | 53 |
| C3 phospho Thr 8 (s8) | 603-611 | 54 |
| C3 c4-Ig like (c4) | 544-633 | 55 |
| C3 linker c4-c5 (l5) | 634-644 | 56 |
| C3 c5-Ig like (c5) | 645-711 | 57 |
| C3 linker c5-f6 (l6) | 712-743 | 58 |
| C3 c6-Fibronectin (f6) | 744-870 | 59 |
| C3 c7-Fibronectin (f7) | 871-967 | 60 |
| C3 linker c7-c8 (l8) | 968-970 | 61 |
| C3 c8-Ig like (c8) | 971-1065 | 62 |
| C3 c9-Fibronectin (f9) | 1066-1163 | 63 |
| C3 linker c9-c10 (lx) | 1164-1180 | 64 |
| C3 c10-Ig like (cx) | 1181-1274 | 65 |

**Figure S1.** Protein domain names, two letter codes (cd), protein sequence assignment (seq), and indexed numbering for domains in the complex βmys/MYBPC3. Domain names begin with M7 or C3 designating origin from βmys or MYBPC3, respectively. Myosin default domains 27k, 50k, and 20k refer to the molecular weights for tryptic proteolytic fragments from cleavage of the MHC sequence in Loop 1 at the active site and Loop 2 in the actin binding site [1]. OM binding (om) is the binding site for Omecamtiv Mecarbil [2-4]. Many domains are identified in the **Figure 3** structures.

| code | phenotypes | index |
|---|---|---|
| d1 | Dilated cardiomyopathy 1A (MyBPC3) | 1 |
| dc | Familial dilated cardiomyopathy | 2 |
| dm | Myopathy, distal, 1 (MYH7) | 3 |
| ds | Dilated cardiomyopathy 1S (MYH7) | 4 |
| h1 | Familial hypertrophic cardiomyopathy 1 (MYH7) | 5 |
| h4 | Familial hypertrophic cardiomyopathy 4 (MYBPC3) | 6 |
| h8 | Familial hypertrophic cardiomyopathy 8 (MYL3) | 7 |
| hb | Hyaline body myopathy (MYH7), Myosin storage myopathy, Myopathy, myosin storage, autosomal recessive | 8 |
| hc | Familial hypertrophic cardiomyopathy, Primary familial hypertrophic cardiomyopathy, Hypertrophic cardiomyopathy, Increased left ventricular wall thickness, Left ventricular hypertrophy, Concentric hypertrophic cardiomyopathy | 9 |
| hd | Cardiomyopathy, hypertrophic, midventricular, digenic (MYLK2) | 10 |
| hx | Familial hypertrophic cardiomyopathy 10 (MYL2) | 11 |
| lc | Left ventricular noncompaction 10 (MYBPC3) | 12 |
| lv | Left ventricular noncompaction cardiomyopathy, Left ventricular noncompaction | 13 |
| lw | Left ventricular noncompaction 5 (MYH7) | 14 |
| m7 | MYH7-Related Disorders | 15 |
| rc | Familial isolated restrictive cardiomyopathy (MYBPC3, MYL3) | 16 |
| uk | conflicting-interpretations-of-phenotype, not specified, uncertain-significance, NA | 17 |

**Figure S2.** Phenotype (ph), 2 letter codes, and descriptive names from the database. Each phenotype is associated with SNVs falling within the genes comprising βmys/MYBPC3 (MYH7, MYL2, MYL3, and MYBPC3). Conflicting-interpretation-of-phenotype (code uk) implies no consensus phenotype from the database. The FHC, DCM, and LVN generalized phenotypes refer to combined sub-phenotypes {hx, h1, h4, h8}, {ds, d1, dc}, and {lc, lv, lw}, respectively.

**NCBI SNP**

Search NCBI SNP database using "GENE missense_variant" search term and capture output in "snp_resultGENE.txt" where GENE={MYH7, MYL2, MYL3 or MYBPC3}.

**Sort**

**Sort** collects SNP rs#'s from "snp_resultGENE.txt" then downloads data from NCBI for each variant and outputs data to file "snp_GENE_missense_variant.dat".

**1. Mlearn**

**M(achine)learn(ing)** initializes 2 or 3 letter codes for population, residue, frequency, phenotype, and pathogenicity characteristics.

**2. SNPs**

**SNPs** extracts data from imported "snp_GENE_missense_variant.dat" file for each SNP characteristic.

**3. Label**

**Labels** assigns domains to SNP substituted residues using sequence location.

**4. DRead**

**DRead** builds matrix containing SNP characteristics. Repeat steps 1-4 for each protein then combine into the complex in a final step.

**Train**

**Train** uses fulfilled 6ddps to train general neural network contraction models in **Figure 1**. They give output (phenotype and pathogenicity) from input (domain, substitution, population, frequency) for unfulfilled 6ddps.

**NnetCyc**

**NnetCyc** computes conditional probabilities (Methods eq 1) from CPTs constructed from 6ddps when unknowns in the data set are given by the general neural network contraction model.

**Wdomain**

**Wdomain** computes joint probability densities (eqs 1-5).

**Lookup Lineup**

**Lookup** computes migrations distances for human populations using two metrics.
**Lineup** indexes migration distances for human populations.

**Correlation**

**Correlation** computes and displays 2D-CG co-domain cross-correlate amplitudes (Methods eqs 6-9).

**Figure S3**. Protocol for SNV data retrieval from the National Center for Biotechnology Information (NCBI). Using a browser, open the NCBI home page, select the SNP database, and use "MYH7 missense_variant" as the search item. The search returns items saved to a text file that is imported to the Mathematica (Wolfram Research, Champaign, IL, USA) program Sort. Sort uses text search/extract tools to identify and collect the SNV reference numbers (rs#) into a list. Redundant rs# entries are merged. Each rs# corresponds to a single reference gene and nucleotide location but may involve various nucleotide substitutions. Mathematica programs in steps 1-4 generally need minor adjustments with each NCBI SNP data build. Other programs appearing in the figure are self explanatory.

| code | population | index |
|---|---|---|
| ACP | ACPOP | 1 |
| AFM | African American | 2 |
| AFR | African | 3 |
| AMR | American | 4 |
| ASI | Asian | 5 |
| ASJ | Ashkenazi Jewish | 6 |
| CEA | CentralAmerican | 7 |
| CRM | Chinese | 8 |
| CUB | Cuban | 9 |
| DAN | Danish | 10 |
| DOM | Dominican | 11 |
| EAS | East Asian | 12 |
| EST | Estonian | 13 |
| EUA | European American | 14 |
| EUR | European | 15 |
| FIN | Finnish from FINRISK project | 16 |
| GON | Genome of the Netherlands | 17 |
| JPN | JAPANESE | 18 |
| KOR | Korean | 19 |
| LA1 | Latin American 1 | 20 |
| LA2 | Latin American 2 | 21 |
| MDE | Middle_Est | 22 |
| MEX | Mexican | 23 |
| NAM | NativeAmerican | 24 |
| NHI | NativeHawaiian | 25 |
| OCE | Oceania | 26 |
| OTH | Other | 27 |
| PCC | PARENT AND CHILD COHORT | 28 |
| PRI | PuertoRican | 29 |
| SAS | SouthAsian | 30 |
| SOA | SouthAmerican | 31 |
| SPC | Spanish controls | 32 |
| TWC | TWIN COHORT | 33 |

**Figure S4**. Human population codes and descriptions from 1000Genomes, gnomAD-Genomes, TopMed, and other studies in the NCBI database including: ACPOP (ACP) whole-genome sequenced control population study from Västerbotten County in northern Sweden; Dominican (DOM) Dominican Republic; Latin American 1 (LA1) Latin American individuals with Afro-

Caribbean ancestry, Latin American 2 (LA2) Latin American individuals with mostly European and Native American Ancestry, Other (OTH) missense SNVs where a population category is not indicated; Parent and Child Cohort (PCC) UK10K Avon Longitudinal Study of Parents and Children Variants; Spanish controls (SPC) Medical Genome Project healthy controls from Spanish population, Twin Cohort (TWC) UK10K Department of Twin Research and Genetic Epidemiology twin registry of 11,000 identical and non-identical twins between the ages of 16 and 85 years.

| Population Sequence | Code | Distance from Addis Ababa km | Line km | Limits km | Outlier | Index |
|---|---|---|---|---|---|---|
| Middle_Est | MDE | 3934 | 3934 | {3518, 5936} | | 1 |
| African | AFR | 4526 | 4526 | {1190, 7002} | | 2 |
| Ashkenazi Jewish | ASJ | 5118 | 5118 | {2480, 8149} | | 3 |
| European | EUR | 5710 | 5710 | {5424, 8149} | | 4 |
| Genome of the Netherlands | GON | 6302 | 6302 | {6000, 9039} | | 5 |
| Danish | DAN | 6894 | 6894 | {6122, 9182} | | 6 |
| PARENT AND CHILD COHORT | PCC | 7486 | 7486 | {6215, 9406} | | 7 |
| European American | EUA | 8078 | 8078 | {6355, 9622} | | 8 |
| SouthAsian | SAS | 8670 | 8670 | {5929, 9815} | | 9 |
| Spanish controls | SPC | 9262 | 9262 | {6703, 10013} | | 10 |
| Asian | ASI | 9854 | 9854 | {6558, 10754} | | 11 |
| American | AMR | 10446 | 10446 | {6687, 10829} | | 12 |
| Oceania | OCE | 11038 | 11038 | {7411, 11531} | | 13 |
| African American | AFM | 11630 | 11630 | {5610, 12193} | | 14 |
| Latin American 1 | LA1 | 12222 | 12222 | {5282, 12662} | | 15 |
| Cuban | CUB | 12768 | 12814 | {7661, 12768} | * | 16 |
| Estonian | EST | 13406 | 13406 | {7862, 14229} | | 17 |
| PuertoRican | PRI | 13998 | 13998 | {8609, 14509} | | 18 |
| Finnish from FINRISK project | FIN | 14590 | 14590 | {7869, 14640} | | 19 |
| East Asian | EAS | 15118 | 15182 | {9572, 15118} | * | 20 |
| Korean | KOR | 15774 | 15774 | {9548, 16416} | | 21 |
| Dominican | DOM | 16365 | 16365 | {10288, 17577} | | 22 |
| SouthAmerican | SOA | 16957 | 16957 | {11171, 18362} | | 23 |
| Latin American 2 | LA2 | 17549 | 17549 | {12425, 19697} | | 24 |
| CentralAmerican | CEA | 18141 | 18141 | {12755, 21166} | | 25 |
| Mexican | MEX | 18733 | 18733 | {13784, 22886} | | 26 |
| NativeAmerican | NAM | 19325 | 19325 | {15912, 28073} | | 27 |
| NativeHawaiian | NHI | 19940 | 19917 | {19940, 31181} | * | 28 |

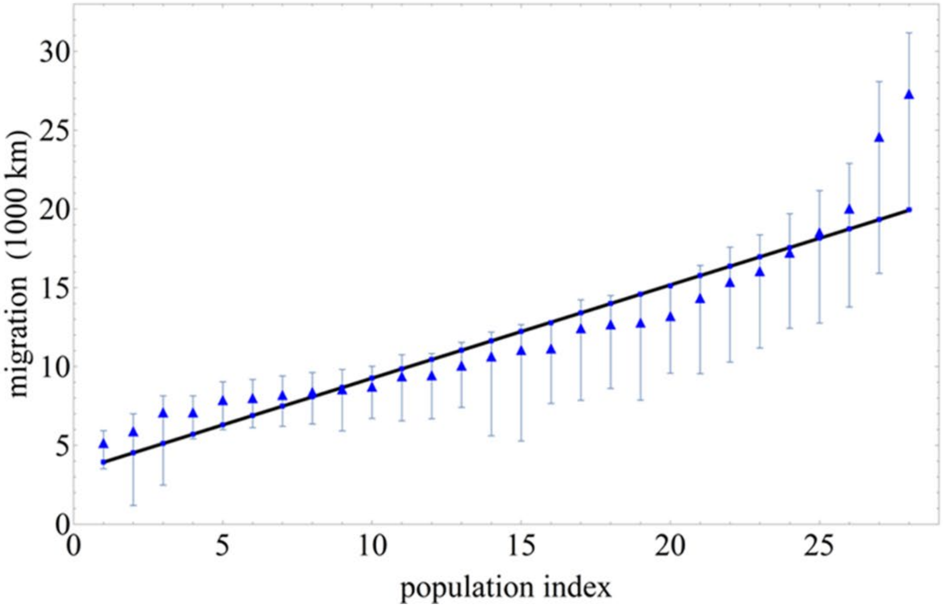


**Figure S5**. Migration distance vs human population index (from **Figure S4**) and pertaining to complex βmys/MYBPC3 in tabular (top) and graphical (bottom) forms. **Top**. Populations (column 1) listed are in a linear relationship with migration distance (column 3). The migration distance is a proxy for genetic divergence variation where divergence decreases with distance. Fitted line (column 4), computed as described [5], closely follows migration distances (column 3) and falls within limits (column 5) except for outliers identified with the asterisk in column 6. The population index (column 7) identifies populations in graphical presentations like that in the bottom panel. **Bottom**. Thick black line indicates a best estimate for the linear relationship of listed populations with migration distance. Light blue vertical bars at each triangle show minimum limits needed to fulfill the linear estimate as described in the text. Blue triangles indicate migration distances falling within the blue vertical bars best fitted by the blue line.

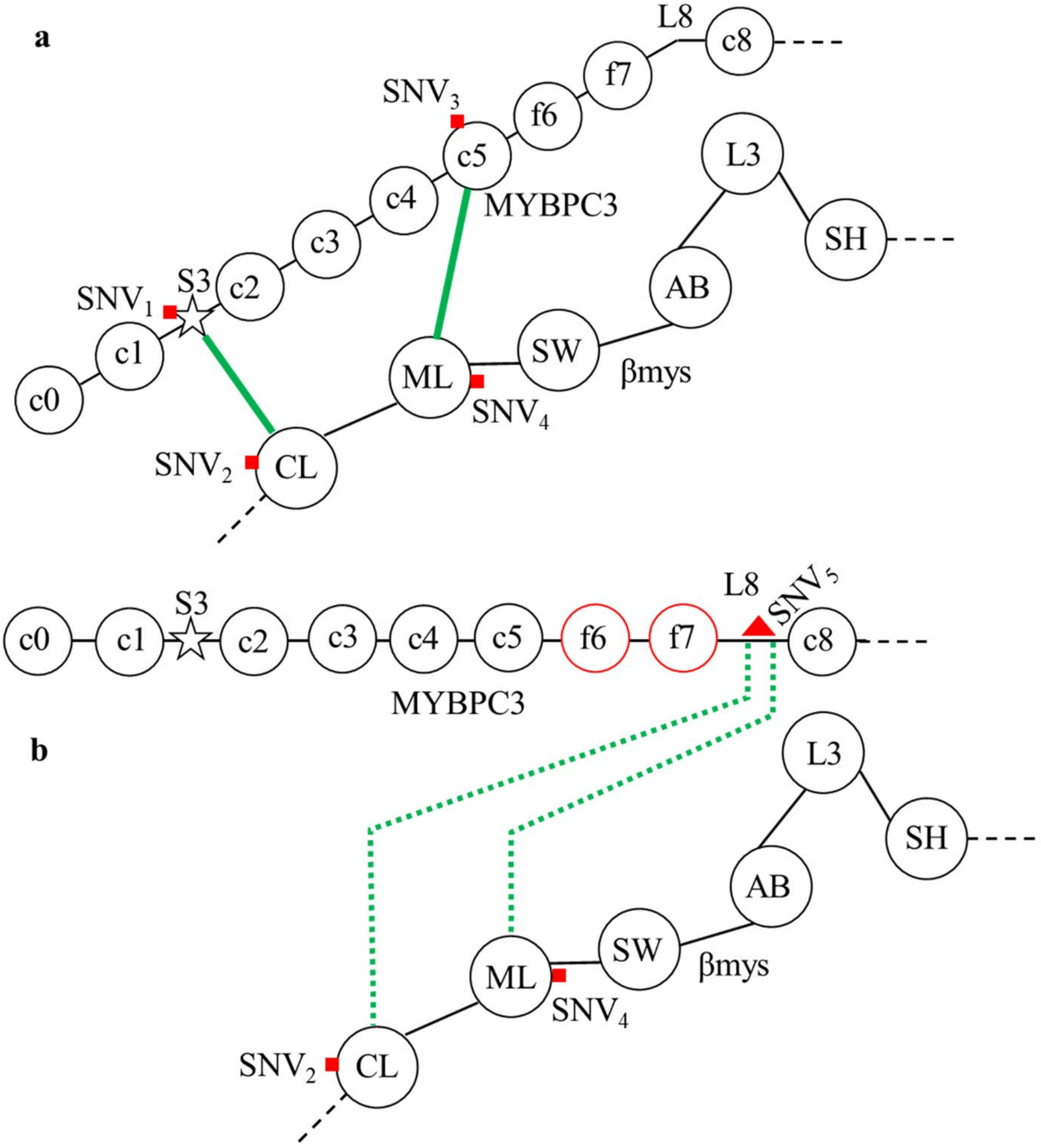


**Figure S6.** Hypothetical βmys/MYBPC3 complexes for human populations with SNVs detecting real (**a**) and virtual co-domains (**b**). (**a**) Functional domains CL and ML in βmys and functional domains S3 (star) and C5 in MYBPC3 have SNVs (red square) detecting the CL-S3 co-domain (by $SNV_1$ and $SNV_2$ probability correlation) and the ML-C5 co-domain (by $SNV_3$ and $SNV_4$

probability correlation). Real co-domains are connected by solid green lines. (**b**) SNV modification of linker 8 (L8) causes conformation change in MYBPC3 and the detection of virtual co-domains CL-L8 (by $SNV_5$ and $SNV_2$ probability correlation) and ML-L8 (by $SNV_5$ and $SNV_4$ probability correlation). Virtual co-domains indicated by dashed green lines. They are virtual because physical contact between the SNV modified domains is unnecessary. Only inter-protein correlates are under consideration.
